# Genome-wide sexually antagonistic variants reveal longstanding constraints on sexual dimorphism in the fruitfly

**DOI:** 10.1101/117176

**Authors:** Filip Ruzicka, Mark S. Hill, Tanya M. Pennell, Ilona Flis, Fiona C. Ingleby, Kevin Fowler, Edward H. Morrow, Max Reuter

## Abstract

The evolution of sexual dimorphism is constrained by a shared genome, leading to ‘sexual antagonism’ where different alleles at given loci are favoured by selection in males and females. Despite its wide taxonomic incidence, we know little about the identity, genomic location and evolutionary dynamics of antagonistic genetic variants. To address these deficits, we use sex-specific fitness data from 202 fully sequenced hemiclonal *D. melanogaster* fly lines to perform a genome-wide association study of sexual antagonism. We identify ~230 chromosomal clusters of candidate antagonistic SNPs. In contradiction to classic theory, we find no clear evidence that the X chromosome is a hotspot for sexually antagonistic variation. Characterising antagonistic SNPs functionally, we find a large excess of missense variants but little enrichment in terms of gene function. We also assess the evolutionary persistence of antagonistic variants by examining extant polymorphism in wild *D. melanogaster* populations. Remarkably, antagonistic variants are associated with multiple signatures of balancing selection across the *D. melanogaster* distribution range, indicating widespread and evolutionarily persistent (>10,000 years) genomic constraints. Based on our results, we propose that antagonistic variation accumulates due to constraints on the resolution of sexual conflict over protein coding sequences, thus contributing to the long-term maintenance of heritable fitness variation.

---

The divergent reproductive roles of males and females favour different phenotypes^1,2^. However, responses to these selective pressures are constrained by a shared genome, leading to ‘sexual antagonism’ where different alleles at given loci are favoured in the two sexes^1,3–5^. A wealth of quantitative genetic studies has established sexual antagonism as near ubiquitous across a wide range of taxa, including mammals^6^ (and humans^7^), birds^8^, reptiles^9^, insects^10,11^, fish^12,13^ and plants^14^. Accordingly, sexual antagonism can be considered a major constraint on adaption and an important mechanism for the maintenance of fitness variation within populations^15^.

However, despite its evolutionary importance, we have little understanding of the biological mechanisms underlying this conflict and virtually no empirical data on the identity and evolutionary dynamics of antagonistic alleles^13^. While a small number of individual antagonistic loci have been identified^12,13^, these are of limited use for elucidating general properties of loci experiencing sexual antagonism. On a genome-wide scale, previous transcriptomic work in *D. melanogaster* has associated antagonistic fitness effects with patterns of gene expression^16^. But despite potentially revealing some of the molecular correlates of fitness variation, this approach cannot distinguish between causal antagonistic loci and their downstream regulatory targets. In humans, genome-wide frequency differences between males and females have been used to infer sexually antagonistic selection on viability^17^, but this approach neglects important reproductive components of fitness. It is essential that we characterise causal antagonistic loci underlying lifetime reproductive success in order to understand the adaptive limits to sexual dimorphism and mechanisms of conflict resolution.

To address this shortcoming, we identified sexually antagonistic loci across the *D. melanogaster* genome and characterised their functional and evolutionary properties. Specifically, we obtained male and female fitness data for over 200 hemiclonal lines that had been extracted from LH_M_, the outbred, laboratory-adapted population in which sexually antagonistic fitness effects were first characterised^10,18^. Our fitness measurements estimate lifetime reproductive success in both sexes by replicating the regime under which LH_M_ has been maintained for over twenty years^19^. We combined these fitness data with high-coverage genome sequences^20^ and performed a genome-wide-association-study (GWAS) to map the genetic basis of sexual antagonism. We then examined the properties of candidate antagonistic polymorphisms, including their genomic distribution across the X chromosome and autosomes, the functional characteristics of candidate polymorphisms and the genes in which they occur, and their population genomic dynamics across a number of wild *D. melanogaster* populations.

## Results

Quantitative genetic analyses confirmed the presence of significant amounts of genetic variation for male and female fitness among the lines assayed (N=223). Estimating the genetic variances and covariances between the lines, we found appreciable heritabilities for fitness in both sexes (female *h*^2^=0.42, 95% CI 0.30–0.54; male *h*^2^=0.16, 95% CI 0.04–0.27). Comparable estimates were also obtained by treating single nucleotide polymorphisms (SNPs) as random effects in a linear mixed model and calculating SNP heritability (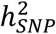) using restricted maximum likelihood (female 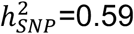, SD 0.13, P<0.001; male 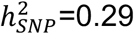, SD 0.16, P=0.007). Overall, the intersexual genetic correlation for fitness did not differ significantly from zero in this sample of genotypes (r_MF_=0.15, 95% CI −0.21–0.46). The presence of ample heritable fitness variation, combined with the lack of a strong positive intersexual genetic correlation for fitness, suggests the presence of sexually antagonistic fitness variants.

We quantified antagonistic fitness variation by calculating an ‘antagonism index’. Specifically, we rotated the coordinate system of the male and female fitness plane by 45 degrees and extracted the position of individual fly lines on the axis ranging from extremely male-beneficial, female-detrimental (MB) fitness effects to extremely female-beneficial, male-detrimental (FB) fitness effects (Fig. 1A). This approach for defining an antagonism index is analogous to other linear transformations, such as the widely applied transformation of human height and weight into a Body Mass Index^21^. The antagonism index itself had high SNP heritability (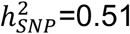, SD 0.15, P=0.001), as expected from the heritability of its sex-specific fitness components.

**Figure 1.**
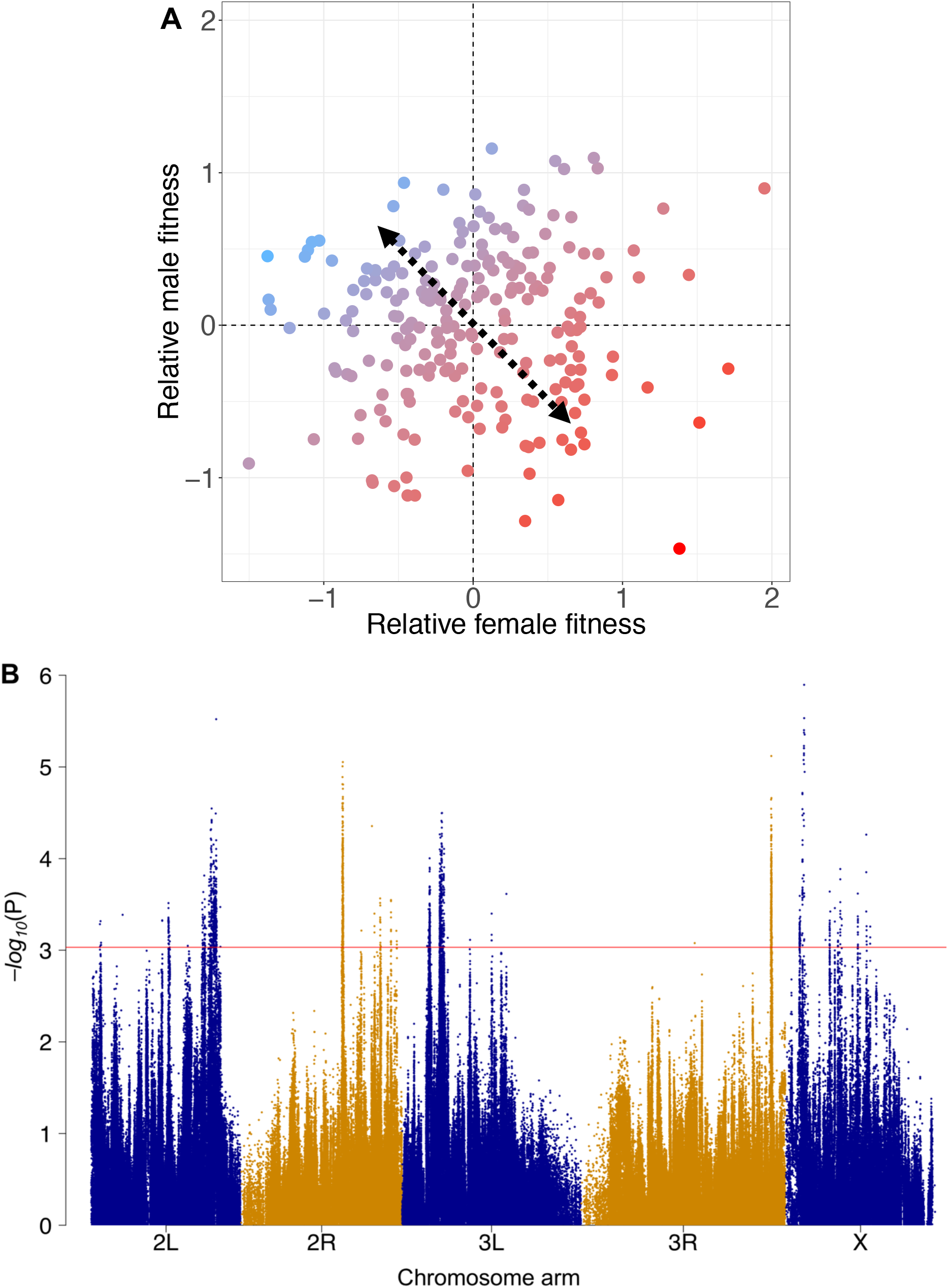
Genome-wide association mapping of sexual antagonism. **A.** Relative male and female lifetime reproductive fitness estimates for 223 *D. melanogaster* hemiclonal lines. Fitness measures have been scaled to be normally distributed. Colours denote each line’s antagonism index, *i.e.* their position along a spectrum (arrows) ranging from male-beneficial, female-detrimental fitness effects (blue), to female-beneficial, male-detrimental effects (red). **B.** Association of each SNP with the antagonism index along the five major *D. melanogaster* chromosome arms, presented as a Manhattan plot where each point represents the −*log_10_* (P) value from a Wald *χ*^2^ association test. Colours denote chromosome arms, the horizontal line represents the Q-value cut-off (0.3) used to define candidate antagonistic SNPs.

To identify putative antagonistic SNPs, we performed a genome-wide association study (GWAS) based on the antagonism index and sequence polymorphism data for 765,654 common (MAF>0.05) and stringently quality-filtered SNPs across 202 of the 223 lines (see Methods; Sup. Fig. 1). We employed a linear mixed model that corrects for between-line relatedness and population structure by incorporating a genetic similarity matrix as a random effect^22^ (Sup. Fig. 2). Figure 1B presents a Manhattan plot of raw P-values from SNP-wise association tests along the *D. melanogaster* genome.

Although no individual SNP reached genome-wide significance based on stringent Bonferroni correction (P<6.53 × 10^−8^), our focus was to characterise broad patterns associated with genome-wide antagonistic variation rather than identifying individual antagonistic sites with high confidence. Accordingly, we applied three main approaches to investigate the general properties of antagonistic SNPs and regions. First, we defined 2,372 candidate antagonistic SNPs (henceforth ‘antagonistic SNPs’) as SNP positions with false discovery rate (FDR) Q-values<0.3 (but note that the Q-values we estimated are likely to be conservative, see Sup. Fig. 2). This threshold achieves a balance between false positives and false negatives that is suitable for genome-wide analyses^23^ and allowed us to contrast the properties of antagonistic and non-antagonistic (Q-value≥0.3) SNPs. Second, we quantified the importance of different classes of SNPs (defined by chromosomal location or function) by partitioning total SNP heritability of the antagonism index (‘antagonistic 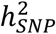’) into the contribution of each class^24,25^. These contributions can then be tested for deviations from random expectations and interpreted without need for defining significance cut-offs for individual SNPs. Finally, we employed set-based association testing where the joint effect of a set of SNPs (such as those in a chromosomal window) on the phenotype is assessed. This joint analysis alleviates the multiple testing burden and can be used to define antagonistic windows with more stringent support (Q-value<0.1). Together these approaches allowed us to characterise the functional properties and evolutionary dynamics of antagonistic genetic variation.

We first examined the genomic distribution of antagonistic variants. The 2,372 antagonistic SNPs were significantly clustered along chromosome arms (median distance: 147bp on autosomes, 298bp on the X chromosome, permutation test: P<0.001 for autosomes and X, Sup. Fig. 3). Using LD clumping^26^, we estimated that the antagonistic SNPs form approximately 226 independent clusters. Some previous theory^3^ and empirical quantitative genetic results^17^ suggest that the X chromosome should harbour a disproportionate amount of antagonistic genetic variation. This was not borne out by our data. We found that, relative to autosomes, the X chromosome neither contained a disproportionate number of antagonistic SNPs (Z-test, P>0.05, Sup. Fig. 4), nor contributed more antagonistic 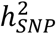 than expected (Fig. 2A).

Our data also allowed us to provide some of the first insights into the biological functions that underlie sexual antagonism. At the most basic level, our results suggest that antagonism arises primarily due to adaptive conflict over coding sequences. Thus, genomic partitioning revealed that variants which result in missense changes were significantly over-represented among antagonistic SNPs (Sup. Fig. 4) and contributed significantly more antagonistic 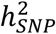 than expected from their proportional genomic representation (Fig. 2A; Sup. Tab. 1). As expected, intergenic regions were under-represented among antagonistic SNPs and contributed qualitatively less antagonistic 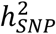 than expected (Fig. 2A; Sup. Fig. 4). However, we found no evidence that SNP functions involved in expression regulation, such as 3’UTR, intronic, upstream or splice region variants, were over-represented among antagonistic SNPs or 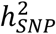 (Fig. 2A; Sup. Fig. 4).

**Figure 2.**
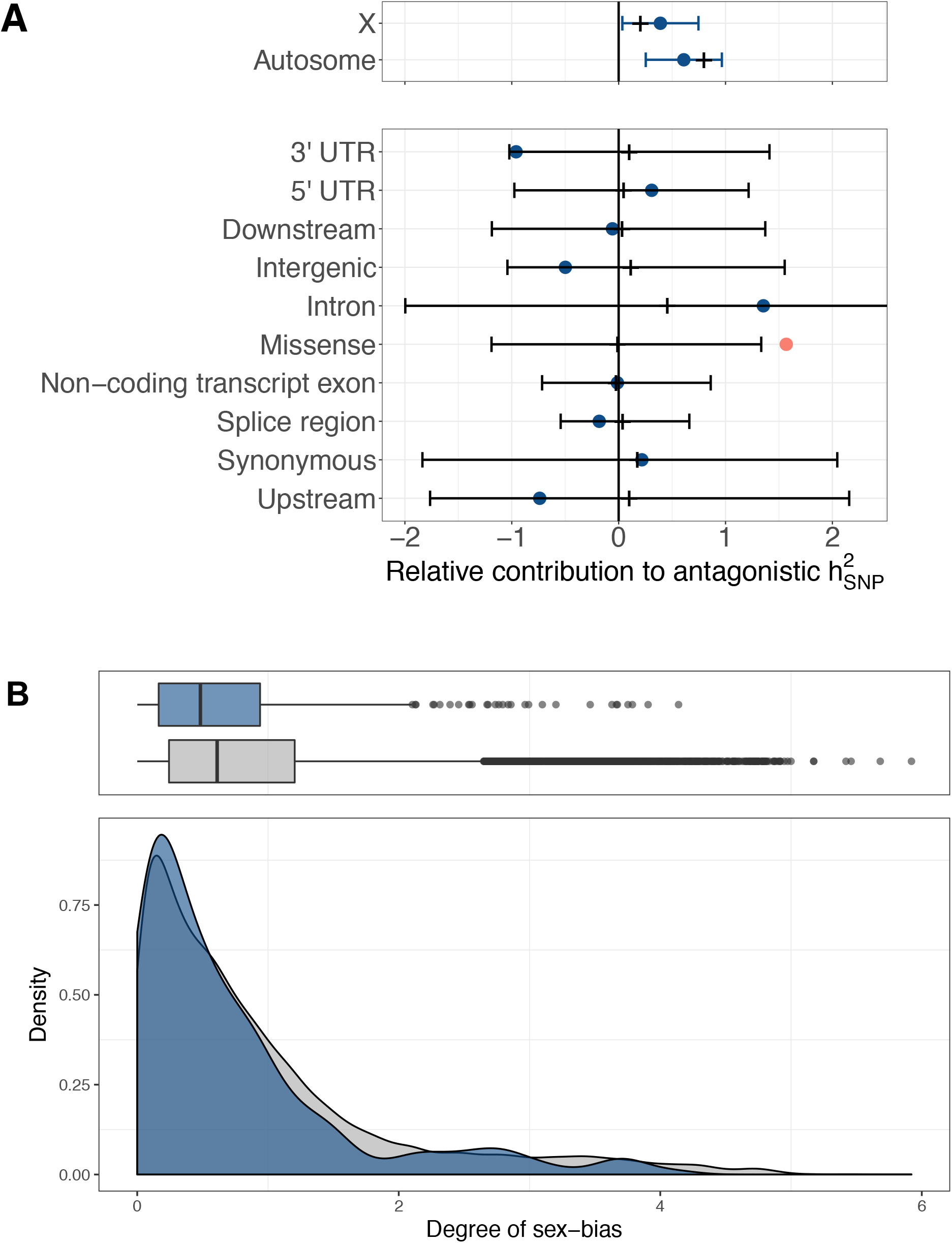
Genomic distribution and functional characteristics of antagonistic variants. **A.** Relative contribution (proportional share) of different chromosomal compartments (top) and functional categories (bottom) to total antagonistic SNP heritability (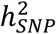). Dots represent estimated 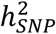 contributions (±95% CI, for chromosomal compartments), with colours indicating significant under or over-representation (red: P-value<0.05; blue: P-value>0.05). Expected 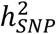 contributions are presented either as black notches (fixed values for chromosomal compartments) or mean±95% CI of the empirical null distribution computed through permutation (for functional categories). See Methods for additional details. **B.** Distributions of the absolute degree of sex-biased expression for antagonistic (blue) and non-antagonistic (grey) genes.

We next performed a series of analyses to characterise the properties of genes harbouring antagonistic SNPs (one or more antagonistic SNPs within ± 5Kb of the gene coordinates). The list of antagonistic genes included some genes known to be involved in sexual differentiation, including *male-specific-lethal-1*, *traffic jam*, and *roundabout 2*, the circadian clock gene *period*, and the Golgi-associated transport protein gene *Tango6* that has been previously found to harbour coding sequence polymorphism shared between *D. melanogaster* and *D. simulans*^27^ (see Sup. Tab. 2 for a complete list of antagonistic genes). Reflecting the heterogeneous list of genes, Gene Ontology (GO) analysis revealed little evidence for preferential association of antagonistic variation with specific biological processes. Only one term, ‘sodium-channel-regulator-activity’, was significant after correction for multiple-testing (Q-value=0.013). However, this annotation is shared by only a few genes (N=5), a cluster of four of which carry antagonistic SNPs. It thus appears that antagonism is not enriched in genes involved in specific functions.

While antagonistic genes were not enriched in specific functions, they did show lower than average sex-bias in gene expression. This pattern is expected because unbiased genes should be most prone to experiencing balanced, opposing selection pressures in the two sexes that would stabilise antagonistic variation. We found evidence for this enrichment both in qualitative terms, where fewer antagonistic genes than expected by chance showed significant sex-biased gene expression (observed=188, expected=212, 11.3% deficit, 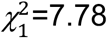, P=0.005), and in quantitative terms, where antagonistic genes had a lower degree of sex-bias than did non-antagonistic genes (W=1309700, P<0.001, Fig. 2B). We did not, however, detect significant overlap between the antagonistic genes identified here and genes that had previously been shown to have sexually antagonistic expression patterns (opposing relationships between expression level and fitness in males and females^16^, observed number of overlapping genes=41, expected number=36, 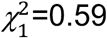, P=0.44). This discrepancy is not unexpected, as the causal genetic changes underlying antagonism need not primarily be associated with expression differences (as suggested by enrichment among missense variants), nor need genes showing expression divergence between the phenotypic fitness extremes necessarily carry causal genetic variants themselves. Finally, we tested whether antagonistic variation is enriched in genes that are likely to be subject to pleiotropic constraints, as this has been proposed to make sexual antagonism harder to resolve^28^. We did not find an association between antagonism and higher levels of pleiotropy, measured either as tissue-specificity^29^ (τ: W=2319200, P=0.80) or as the number of protein-protein interactions^30^ (PPIs: F_1,5276_=2.43, P=0.12). This implies that pleiotropy—at least as captured by τ and PPIs—does not contribute significantly to maintaining sexually antagonistic genetic variation.

In addition to assessing the functional properties of antagonistic loci, we also investigated the population genetic effects of sexual antagonism. Models predict that the opposing sex-specific fitness effects of antagonistic alleles generate balancing selection, resulting in elevated levels of genetic polymorphism at antagonistic loci^31–33^. Having identified putatively antagonistic variants, we can test this prediction by comparing levels of polymorphism at antagonistic and non-antagonistic loci. Doing so directly within the LH_M_ population is problematic, because the power to detect antagonistic effects is higher at more polymorphic sites, and candidates therefore tend to show above-average polymorphism. However, we can use data from independent populations and ask whether, for a given level of polymorphism in LH_M_, polymorphism there is greater at antagonistic than at non-antagonistic sites. We performed this type of analysis (see Methods for details) using publicly available polymorphism data from the *Drosophila* Genetics Reference Panel^34,35^ (DGRP), a collection of 205 wild-derived inbred lines. Like LH_M_, the DGRP was established from a North American *D. melanogaster* population. Given the relatively recent colonisation of the continent by *D. melanogaster* (~150 years^36^), the two populations are closely related. We found that antagonistic SNPs had elevated minor allele frequencies (MAFs) in the DGRP, although owing to the close relationship between LH_M_ and the DGRP (and the resulting similarity in allele frequencies) this difference was not statistically significant (P=0.322, Fig. 3A,B). However, when using the P-value for the antagonistic effects of individual SNPs rather than a binary antagonistic/non-antagonistic categorisation of sites, we found a significant negative correlation between P-value and MAF in the DGRP, consistent with elevated polymorphism in the DGRP at SNPs that are more closely associated with antagonism in LH_M_ (*ρ*=−0.055, P=0.044, Fig. 3C). This evidence for antagonism-driven balancing selection at individual sites was corroborated by patterns of regional polymorphism—measured as Tajima’s D within 1000bp windows along the chromosome arms. Tajima’s D was significantly higher in antagonistic windows (those with Q-value<0.1 in a window-based GWAS) than in non-antagonistic windows (Q-value≥0.1; F_1,115477_=224.6, P<0.001, Fig. 3G). Overall, these analyses show that the heritable phenotypic variation in sex-specific fitness that can be generated and maintained by sexual antagonism is mirrored by a signal of increased polymorphism at the underlying genetic loci.

**Figure 3.**
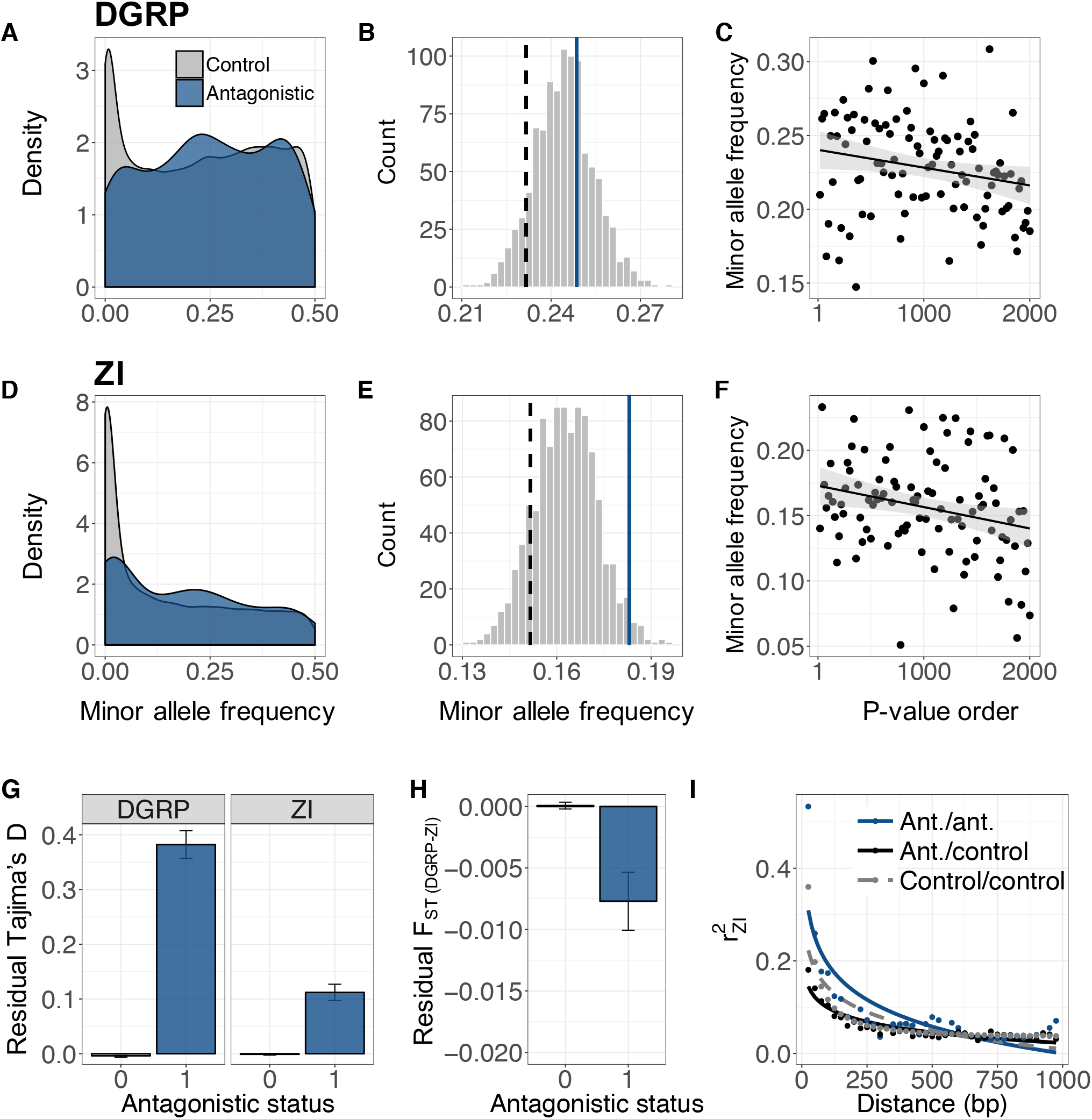
Signatures of balancing selection associated with antagonistic variants in two independent populations (DGRP and ZI; see Sup. Fig. 5 for SA population). **A,D.** Spectra of raw minor allele frequencies (MAF) for LD-pruned antagonistic (blue) and non-antagonistic (‘control’, grey) SNPs in the DGRP and ZI populations. **B,E.** Distribution of mean MAFs for 1,000 sets of LD-independent, non-antagonistic SNPs that have been frequency-matched to LH_M_ antagonistic SNPs (see Methods). Blue line denotes mean MAF of antagonistic SNPs; black dashed line denotes mean MAF of non-antagonistic SNPs before frequency-matching. **C,F.** MAF in the DGRP and ZI populations across 100 sets LD-independent SNPs, each set matched for LH_M_ allele frequencies, and presented in ascending order by P-value. For visualisation purposes, a linear regression line (±95% CI) is shown. **G.** Mean (±S.E.) residual Tajima’s D (corrected for linked selection, see Methods) for antagonistic windows (blue; ‘antagonistic status=1’) and non-antagonistic windows (grey; ‘antagonistic status=0’) in the DGRP and ZI populations. **H.** Residual F_ST_ (±S.E.), corrected for linked selection (see Methods), for antagonistic and non-antagonistic windows in the DGRP and ZI populations. **I.** Linkage disequilibrium (r^2^) in the ZI population between pairs of antagonistic SNPs (blue, ‘Ant./ant.’), pairs of non-antagonistic SNPs (grey, ‘Control/control’) and mixed pairs (black, ‘Ant./control’). Points represent mean r^2^ across 25bp bins; r^2^ is modelled as a declining exponential function of distance (fitted lines).

A key, yet so far unresolved question is whether antagonistic polymorphisms are mainly short-lived and population-specific or persist over prolonged periods of time. The analyses of polymorphism in the DGRP shed some light on this question, demonstrating that antagonistic polymorphisms are maintained at least over periods of tens to hundreds of years, or hundreds to a few thousand generations. In order to assess signals of balancing selection over longer time spans, we repeated these analyses with data from a population in *D. melanogaster*’s ancestral Sub-Saharan distribution range, in Zambia (ZI, 197 genomes from phase 3 of the *Drosophila* Population Genomics Project^37^; see also Sup. Fig. 5, which repeats the ZI analyses with identical results using 118 genomes from South Africa^38^). Just as in the DGRP, we found that antagonism generated a clear signature of balancing selection in this ancestral population sample. Analyses based on binary categories showed that antagonistic SNPs had significantly higher MAFs in ZI compared to non-antagonistic SNPs (P=0.024, Fig. 3D,E), while analyses based on P-values showed again that sites with stronger evidence for antagonistic effects had more elevated MAFs (*ρ*=−0.070, P=0.002, Fig. 3F). At a larger chromosomal scale, antagonistic windows had significantly higher polymorphism (Tajima’s D) than non-antagonistic windows (F_1,116099_=60.63, P<0.001, Fig. 3G). Furthermore, they also exhibited lower population differentiation between DGRP and ZI (measured as F_ST_; Wilcoxon Rank-Sum test, W=63416000, P=0.012, Fig. 3H; Sup. Fig. 5), in line with balancing selection maintaining similar frequencies across distant populations.

In addition to elevated polymorphism in antagonistic regions of the genome, we also found evidence for increased linkage disequilibrium (LD) – another hallmark of balancing selection^39^. We compared local LD (<1,000bp, measured as r^2^) between pairs of antagonistic sites, pairs of non-antagonistic sites, and ‘mixed’ site pairs (consisting of an antagonistic and a non-antagonistic SNP) in the ZI population, which is most phylogenetically distant from LH_M_ and where a signal of LD should be weakest in the absence of long-term balancing selection. Consistent with selection, we found that pairs of antagonistic sites had higher LD in this population than pairs of non-antagonistic sites (Wilcoxon Rank-Sum tests, W=8346500000, P<0.001, Fig. 3I). They also had higher LD relative to mixed pairs (Wilcoxon Rank-Sum test, W=33823000, P<0.001, Fig. 3I). Thus, high LD between antagonistic sites is not an artefact of unusually low levels of recombination near antagonistic regions, but instead reflects the action of long-term balancing selection.

Taken together, these comparative population genomic analyses demonstrate that the antagonistic allelic variation identified in LH_M_ is neither recent nor population-specific. To a significant degree, balancing selection maintains antagonistic variation over timescales that extend beyond the extension of the species range out of Africa, more than 10,000 years ago^36^.

## Discussion

Our study provides the first genome-wide analysis of the identity, function and evolution of sexually antagonistic sequence polymorphisms in fruitflies. Remarkably, we find that genetic variation at antagonistic loci is stably maintained across *D. melanogaster* populations throughout the species’ distribution range, indicating that the targets of antagonistic selection have been largely conserved for many millennia^36,40–42^—and several tens of thousands of generations. The geographical stability and low turnover in antagonistic sequence variation implies that adaptive conflict between males and females is rooted in a fundamental aspect of the biology of the sexes and persists even in the face of environmental variation^43^. It is therefore unaffected by the adaptation of populations to the environmental conditions that they encountered during their colonisation of the globe^36,41,44^ or the continuous adaptive evolution that occurs within temperate populations over the course of the seasons^23^. More generally, our results supplement a growing body of evidence^23^ suggesting that balancing selection can influence patterns of genetic variation on a genome-wide scale, rather than simply a small number of isolated loci^45^, as is often assumed^46^. Sexually antagonistic selection should contribute particularly strongly to the build-up of balanced polymorphisms, given that there is abundant evidence for sex-specific selection in nature^4,47^ and that sex-specific selection can generate permissive conditions for the evolution of such polymorphisms relative to alternative modes of balancing selection^33,48^.

The long-term stability of sexually antagonistic polymorphisms further suggests that the evolutionary constraints on sexual dimorphism inherent in antagonism are difficult to resolve. While we do not find any evidence that there is elevated pleiotropy among genes experiencing ongoing conflict, the persistence of antagonism fits with our finding that antagonistic polymorphisms are highly enriched for missense variants. While antagonistic selection on expression levels can be accommodated by gradual evolution of sex-specific gene expression^49^, adaptive conflicts over coding sequences can only be resolved through a complex multi-step process^50^ of gene duplication, sex-specific sub-functionalisation of coding sequences and the evolution of differential expression of the two paralogues. The requirement for gene duplication, in particular, would be expected to constitute a severely limiting barrier for this route towards resolution, as suitable mutation events will be exceedingly rare. This large barrier to resolution, and the resulting stochasticity in which antagonisms will undergo resolution, may help to explain the lack of GO enrichment observed among antagonistic genes.

We find no convincing evidence that the X chromosome is enriched for antagonistic variation, in contradiction to classic theory^3^. This discrepancy could be due to the presence of dominance reversal, where the beneficial allele is dominant in each sex. Such sex specific dominance has recently been documented empirically^13^ and is predicted to shift enrichment of antagonism from the X to the autosomes^51^—particularly so if antagonistic loci interact epistatically^52^. Furthermore, the hemiclonal approach might miss low-frequency X-linked antagonistic polymorphisms with recessive fitness effects, as these will rarely be expressed in phenotypic assays. However, while these general effects might explain the lack of X-enrichment, our result also contradicts previous empirical findings obtained in the LH_M_ population^17^ itself, which found that the X chromosome contributed disproportionally to antagonistic fitness variation. The previous study was based on a much smaller sample of genomes, with large uncertainties about the estimated chromosomal contributions. It was also performed more than ten years ago and much closer to the establishment of LH_M_ as a laboratory population. Accordingly, the discrepancy to our results might in part be explained by stronger genetic drift on the X chromosome relative to autosomes, which could in turn lead to a disproportionate loss of X-linked antagonistic polymorphisms^32^.

Taken together, this study addresses a longstanding gap in our understanding of sexual antagonism, and provides a valuable resource from which to further elucidate the origin and resolution of this fundamental evolutionary phenomenon.

## Methods

### LH_M_ hemiclones

LH_M_ is a laboratory-adapted population of *Drosophila melanogaster* that has been maintained under a highly controlled rearing regime since 1996^53^. A random sample of 223 genetic lines was created from the population^20^ using a hemiclonal approach^54^. Individuals of each line carry an identical haploid genome comprising the major chromosomes X, 2 and 3. Crosses with flies from custom stocks allows the generation of many replicate individuals—males and females—that carry a line’s X-2-3 haplotype alongside a random chromosomal complement from the LH_M_ population that can be assayed for fitness.

### Fitness measurements

Lifetime adult reproductive fitness of males and females of each line was measured using assays designed to mimic the LH_M_ rearing regime. For male fitness, we measured competitive fertilisation success by setting up competition vials containing 5 hemiclonal males from a given line, 10 competitor *bw* males and 15 virgin *bw* females. After two days, *bw* females were isolated into individual vials containing no additional yeast and left to oviposit for 18 hours. On day 12 post egg-laying, progeny were scored for eye colour. Male fitness was calculated as the proportion of offspring sired by the 5 hemiclonal males (those with wildtype eye-colour), combining progeny data from the 15 oviposition vials. This assay was repeated 5 times in a blocked design; estimates for each line were therefore based on fitness measurements from 25 hemiclonal males.

Female fitness was measured as competitive fecundity. Competition vials containing 5 virgin hemiclonal females from a given line, 10 competitor *bw* females and 15 *bw* males were set up. Two days later, the 5 hemiclonal females were isolated into individual vials and left to oviposit for 18 hours. These vials were immediately chilled at 4°C and fecundity was measured by counting the number of eggs laid per female. This assay was replicated 5 times in a blocked design; each line estimate therefore measured the fitness of 25 hemiclonal females.

Fitness data were subjected to quality control and pre-processing in preparation for quantitative genetic and association analysis. Male fitness data from competition vials where not all 5 focal males were present at the end of the assay were removed from further analysis. Similarly, we omitted female oviposition vials where fewer than 2 eggs were present (indicating partial sterility or failure to mate) or where the female had died over the course of the assay. For each sex, fitness measurements were then first box-cox transformed to be normally distributed within each block, then scaled and centred. To calculate SNP heritabilities and for association analysis, data from each block were averaged to obtain one fitness estimate for each line and sex.

### Quality control of whole-genome sequences

We used previously published whole genome sequences generated from the hemiclonal lines analysed here^20^. Details about DNA extraction, library preparation, sequencing, read processing and SNP calling are provided in the original publication. Prior to the association analysis performed here, further site-level quality filtering steps were performed in vcftools^55^ and *PLINK*^26^. First, individual variant calls based on depth<10 and genotype quality<30 were removed. Second, individuals with>15% missing positions were removed. Third, positions with poor genotype information (<95% call rate) across all retained individuals were discarded. Finally, given the relatively small sample size of the dataset as a whole and the low power of an association test for rare variants, we retained only common variants (MAF>0.05) for further analysis. From an initial dataset of 220 hemiclones containing 1,312,336 SNPs, this yielded a quality-filtered dataset of 765,980 SNPs from 203 hemiclones.

To detect outliers, we examined LH_M_’s population structure using principal components analysis (PCA). Overlapping SNP positions from the 203 LH_M_ genomes and from an outgroup population (*Drosophila* Genetic Reference Panel, or DGRP^34^) consisting of 205 whole-genome sequenced individuals were used as input to construct a genetic similarity matrix. This set of SNPs was pruned for linkage disequilibrium (LD) such that no two SNPs with r^2^>0.2 within 10Kb remained. The leading PC axes were extracted in LDAK (“Linkage-Disequilibrium Adjusted Kinships”^56^). After removal of one outlier (see Sup. Fig. 1A), the final dataset used for association analysis contained 202 individuals and 765,764 SNPs.

### Heritability analyses

We estimated the variance-covariance matrix for fitness and sex-specific residual variances by fitting a model using MCMCglmm^57^ implemented in R. Specifically, we fitted the model *Y_ijk_* = *X_ij_* + *ε_ijk_*, where *Y_ijk_* is the scaled and centred fitness of individual k of genotype j and sex i, *X_ij_* is the sex-specific random effect of genotype j in sex i, and *ε_ijk_* describes the sex-, genotype- and individual-specific residual. The genotypic fitness effects in males and females follow a bivariate normal distribution *X_ij_*~*N*(0,G), where

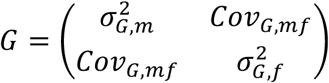

is the genetic variance-covariance matrix across sexes (composed of male and female additive genetic variances 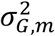 and 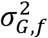 and the intersexual genetic covariance *Cov*_G,mf_). Residuals follow a normal distribution 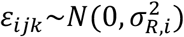, where 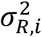 is the sex-specific residual variance, and are assumed to be uncorrelated across sexes.

From these variance estimates, we calculated male and female heritabilities of fitness as 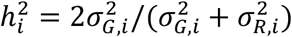, where the subscript i indicates either male or female. The factor 2 in the heritability calculation reflects the fact that with the hemiclonal approach, individuals assayed share half their genetic material (the hemizygous hemiclonal genome). The intersexual genetic correlation was calculated as 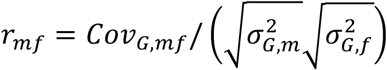. The quantitative genetic parameters 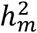, 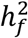 and *r_mf_* were calculated for each sample from the Monte Carlo Markov chain. From these series of values we obtained point estimates (averages) and 95% credible interval (using the function HPDintervals).

As a complementary approach, we estimated the SNP heritability (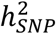) of male and female fitness in LDAK^56^. This approach uses Restricted Maximum Likelihood (REML)^58^ to fit a linear mixed model that expresses the vector of phenotypes Y as a function genome-wide SNP genotypes, treated as random effects:

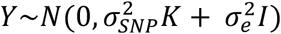

where K the kinship matrix, 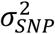 a vector of additive genetic variances for each SNP, 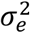 the vector of residual variances and I an individual identity matrix. SNP heritability is then estimated as 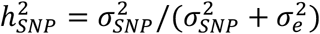.

LDAK corrects for local linkage when calculating SNP heritabilities to avoid inflation of 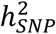 in clusters of linked sites that otherwise arises because several SNPs tag the same causal polymorphism. SNPs are weighted inversely proportional to their local linkage, such that SNPs in high LD contribute less to 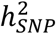 than SNPs in low LD. This model has been shown to substantially improve heritability estimates across a wide range of traits^24^. LDAK also allows to set the parameter α that determines how SNPs are weighted by their MAF (as MAF^α^) when calculating the kinship matrix K. We used the default of α=−0.25 which provides a steeper relationship between MAF and 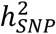 than the value of −1 that is frequently used in studies on humans. Significance of 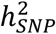 estimates was assessed by permuting phenotype labels 1,000 times, re-calculating 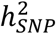 on each permutation as above, and calculating the number of permuted estimates which exceeded the observed.

### Quantification and association analysis of sexual antagonism

To identify loci underlying sexual antagonism, we defined an antagonism index (see main text, Fig. 1A). We calculated its SNP heritability (‘antagonistic 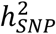’) in LDAK, following the same procedure and settings as those for estimating sex-specific SNP heritabilities.

We performed a GWAS by applying a linear mixed model to test the effect of allelic variants at each SNP on the antagonism index, while including the kinship matrix as a random effect to account for the heritable portion of genetic variation attributable to kinship between individuals. This approach has been shown to effectively control the false positive rate and increase power to detect true associations in samples with moderate degrees of population structure and close relatedness, such as LH_M_^22,59^. The GWAS was implemented in LDAK (settings as above) and a Wald *χ*^2^ test was used to generate P-values for each position.

The genomic inflation factor^60^ of *λ*_median_=0.967 (calculated as median *χ*^2^_obs_/median *χ*^2^_exp_ in GenABEL^61^ in R) suggests that genetic confounding has been well controlled in our GWAS. This is corroborated by the fact that the distribution of P-values when permuting the phenotype labels (100 times and applying the same linear mixed model) was not enriched for low P-values (such a pattern would be expected if residual genetic confounding remained in our sample).

### Defining candidate antagonistic SNPs and regions

We corrected for multiple testing using a False Discovery Rate (FDR) approach and converted P-values into Q-values. We defined antagonistic SNPs as sites with FDR Q-values<0.3 and non-antagonistic SNPs as sites with Q-value≥0.3.

For analyses which consider larger genomic regions (windows), we ran a set-based association test implemented in LDAK (options using ‘–calc-genes-reml’, ‘ignore-weights YES’ and α=−0.25). The test calculates set-wide 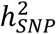 via REML, corrects for local relatedness using the predictors in each window, and computes a P-value using a likelihood ratio test (LRT). The sets we used were 1000bp windows (500bp step) defined according to *Drosophila* Reference 5 genome coordinates, and subsequently converted (using UCSC’s liftOver tool^62^) to Release 6 coordinates. This was a necessary step, as publicly available polymorphism data was mapped to Release 5 of the *D. melanogaster* genome, whereas the GWAS data was mapped to Release 6. We then calculated window-based Q-values from the LRT P-values and defined antagonistic windows as those with Q-value<0.1.

### Genomic distribution of antagonistic SNPs

To estimate the number of independent antagonistic regions, we performed LD-clumping in *PLINK*. We used a significance threshold of 0.00093 for the index SNP (the maximum, least significant, P-value across all antagonistic SNPs), and clustered (‘clumped’) neighbouring antagonistic SNPs by specifying an r^2^ threshold of 0.4 and a distance threshold of 10Kb.

We also quantified the clustering by calculating the median distance between all pairs of adjacent antagonistic SNPs across chromosome arms. We did this separately for the autosomes and X chromosome, to accommodate for the lower SNP density on the X chromosome. We tested for significant clustering by using a permutation test, where antagonistic/non-antagonistic labels was permuted among all SNPs, distances between adjacent SNPs labelled as ‘antagonistic’ after permutation were recalculated as before and the median distance recorded. This process was repeated 1,000 times in order to generate a null distribution of median distances. Significance of clustering among true antagonistic SNPs was calculated as the proportion of median distances in the null distribution that were lower than or equal to the true median distance.

To examine the proportional contribution of autosomal and X-linked antagonistic variants to total 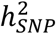, we used two complementary methods. First, we partitioned the genome into X chromosome and autosome subsets, and calculated 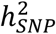 via REML in LDAK each subset in turn (default parameters; α=−0.25). The observed proportion of 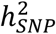 contributed by each compartment was then compared to the expected proportion (*i.e.*, the fraction of LD-weighted predictors belonging to each compartment). We tested whether the two compartments contributed significantly more 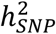 than expected using a two-sample Z-test. Second, we compared the proportion of antagonistic SNPs (Q-value<0.3) to the proportion of all SNPs mapping to each chromosomal compartment, using Z-tests. The under- or over-representation of antagonistic SNPs (deficit or excess of antagonistic compared to all SNPs) in each compartment is therefore unaffected by differences in SNP density between chromosome arms, such as the lower density on the X chromosome.

### Functional analyses of antagonistic loci

We used the variant effect predictor (Ensembl VEP^63^) to map SNPs to functional categories. We partitioned total antagonistic 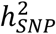 into functional subsets, and estimated the observed proportion of 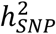 contributed by each subset using REML in LDAK (settings as above). We then used a permutation test to compare observed and expected 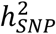 for each functional category, where we shifted genome-wide annotations to a random starting point along a ‘circular genome’. This procedure breaks the relationship between each SNP and its annotation while preserving the order of annotations and their associated LD structure^64^. 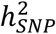 was re-calculated via REML for each of 1,000 permuted datasets and two-tailed P-values determined as the sum of permuted estimates with more extreme absolute values than the observed. As a complementary approach, we compared the proportion of antagonistic SNPs to the proportion of all SNPs mapping to each functional category. We then assessed enrichment for each functional category in turn using Z-tests.

We also used the VEP to map SNPs to genes. We included extended gene regions (+/- 5kb of gene coordinates, VEP default) in our gene definition. To gain preliminary insights into the functions of antagonistic genes we used the Gorilla^65^ Gene Ontology tool, with FDR correction for multiple testing across GO terms. All genes covered in the final SNP dataset were used as the background set.

To examine the relationship between antagonistic genes and sex-biased gene expression we used the Sebida online database^66^ to annotate genes as having either sex-biased or unbiased expression profiles (meta-class identifier). We then used a *χ*^2^ test to compare the sex-biased expression status of antagonistic and non-antagonistic genes. We additionally examined the quantitative degree of sex-bias using this same dataset. We took the absolute value of the log2 transformed ‘M_F’ bias variable, such that large values indicate more extreme sex bias in expression irrespectively of whether this bias is towards males or females. We compared the distributions of this variable between antagonistic and non-antagonistic genes using a Wilcoxon Rank-Sum test.

To assess the degree of overlap between antagonistic genes identified here and those associated with sexually antagonistic expression patterns in a previous study^16^, we included only genes covered in both datasets, and only those genes in both datasets that were adult-expressed. To determine whether genes were adult-expressed we used the *Drosophila* gene expression atlas (FlyAtlas^67^). Conservatively, we considered a gene ‘adult-expressed’ if its transcript was detected as present in at least one library of one adult-derived sample. We then used a *χ*^2^ test to assess the degree of overlap between the datasets.

We used the tissue-specificity index (τ) to compare pleiotropy between antagonistic and non-antagonistic genes. We used gene expression data from FlyAtlas^67^ to get average expression values for each gene and in each tissue and then calculated τ as:

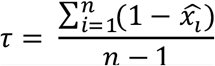

where 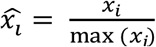 is the proportional expression level of the gene in tissue *i* and *n* is the number of tissues. We compared values of τ for antagonistic and non-antagonist genes using a Wilcoxon rank-sum test.

As an additional proxy for pleiotropy we examined the number of protein-protein interactions (PPIs) between antagonistic and non-antagonistic genes. We used the physical interactions table from FlyBase^68^ to summarise the total number of PPIs for all genes and then compared candidate and non-candidate genes using a general linear model (GLM) with quasipoisson error structure to account for overdispersion.

### Comparative population genomic data

To analyse SNP polymorphism outside the LH_M_ population, we used publicly available population genomic data from three wild *D. melanogaster* populations. The first is an introduced population from North America (*Drosophila* Genetic Reference Panel^34,35^, denoted DGRP: 205 whole-genome sequences derived from inbred lines). The two others come from *D. melanogaster*’s ancestral distribution range in sub-Saharan Africa (Zambia–ZI: 197 whole-genome sequences derived from haploid embryos; South African–SA: 118 whole-genome sequences derived from inbred lines, combines data from sub-populations ‘SD’ and ‘SP’, which have very low population differentiation^38^).

All genome sequences were downloaded as FASTA files from the Drosophila Genome Nexus website (www.johnpool.net/genomes.html). These files had been generated following standardised alignment and quality filtering steps^37^ and were further quality-filtered for admixture and identity-by-descent using scripts provided on the Genome Nexus website. We used snp-sites^69^ to call SNPs and convert the multiple sequence alignments to vcf format. Allele frequencies in the three populations were calculated using vcftools. We further excluded tri-allelic and poorly covered sites (call rate<20).

### SNP-based analyses of balancing selection

To test whether antagonistic sites are associated with signatures of balancing selection, we closely followed the approach of Turchin et al.^70^ and looked for an increased minor allele frequency (MAF) at antagonistic relative to non-antagonistic sites (as identified in LH_M_) in the three comparison populations (DGRP, ZI, SA). By focussing on the contrast between classes of SNPs, we ensured that demographic differences between populations did not confound our analyses.

We first LD-pruned the LH_M_ dataset by clumping (in *PLINK*) to avoid pseudo-replication due to correlations between SNPs. For antagonistic sites, we used the 226 index SNPs identified in the previous clumping. For non-antagonistic sites, a non-antagonistic SNP was randomly chosen as an index SNP and clumped by clustering all SNPs within 10kb with r^2^>0.4. Pruning in this manner reduced the original dataset of 765,764 SNPs to 36,319 “LD-independent” SNPs. For each of these SNPs, we then estimated MAFs in each comparison population. We assigned MAF=0 to sites which were monomorphic in a comparison population and those where a comparison population was polymorphic for variants other than those segregating at that site in the LH_M_.

We then used this LD-independent dataset to compare MAF between antagonistic and non-antagonistic SNPs. We did this using a Monte Carlo approach where, 1,000 times, we paired the 226 antagonistic SNPs with 226 randomly drawn non-antagonistic “control” SNPs. The latter were carefully frequency-matched to the 226 antagonistic SNPs. The matching procedure first corrected LH_M_ MAF for ‘linked selection’^71^ by taking the residuals of a linear regression of LH_M_ MAF on estimates of linked selection. These estimates quantify local recombination rates and proximity to functional sequences in *D. melanogaster*. They thereby account for factors that affect polymorphism along the genome, such as background selection and selective sweeps. We then drew sets of 226 non-antagonistic SNPs to match the residual LH_M_ MAF distribution of the 226 antagonistic SNPs and for each set calculated the mean MAF in the comparison population. The 1,000 sets generated in this way provided a null distribution of MAFs for non-antagonistic sites in each comparison population. P-values for deviations in polymorphism between antagonistic and non-antagonistic sites were then calculated by comparing, in each population, the mean MAF of the 226 antagonistic SNPs to the null MAF distribution.

A second analysis used the same LD-independent dataset but considered the whole spectrum of P-values, rather than a binary split of SNPs into antagonistic/non-antagonistic categories. To this end, we performed binning in two dimensions, by residual LH_M_ MAF (20 quantiles) and P-values (100 quantiles). We then drew one SNP from each of these MAF/P-value bins (2,000 SNPs in total), recorded the MAF for each in the comparison population of interest, and finally correlated these MAF values with P-values of the associated SNPs in the LH_M_ using a Spearman’s rank correlation. Under the hypothesis of antagonism-mediated balancing selection, SNPs with low P-values should tend to have higher MAFs in the population under consideration than SNPs with high P-values.

### Window-based analyses of balancing selection

We performed genome-wide sliding window analyses (1,000bp windows, 500bp step size) to investigate regional signatures of balancing selection. Tajima’s D, which compares SNP polymorphism (nucleotide diversity, π) to SNP abundance (Watterson’s estimator, θ_W_), was compared for windows defined as antagonistic (Q-value<0.1) or non-antagonistic (Q-value≥0.1) from the set-based analysis (see section ‘Defining candidate antagonistic SNPs and regions’). Under the hypothesis that antagonism generates balancing selection, Tajima’s D is expected to be elevated in antagonistic windows. We calculated Tajima’s D for each comparison population using PopGenome in R^72^. As in SNP-based analyses, we incorporated estimates of linked selection^71^ (estimated in 1,000bp windows) by calculating the residuals of a regression of Tajima’s D on estimates of linked selection. Since estimates of linked selection were not available for windows on the X chromosome, we instead used estimates of recombination rate on this chromosome^73^. We then used a generalised linear model (GLM), assuming Gaussian error structure, to compare residual Tajima’s D between antagonistic and non-antagonistic windows.

We also tested for another signature of balancing selection, reduced population differentiation. Measures such as F_ST_ are often considered problematic because they do not correct for the dependency of F_ST_ on local levels of polymorphism^74^. However, the availability of genome-wide estimates of linked selection in *D. melanogaster* allowed us to incorporate this confounding variable explicitly. We therefore estimated F_ST_ over windows, using PopGenome, correcting F_ST_ for linked selection in a way analogous to that used for Tajima’s D. Since the distribution of F_ST_ values is not normally distributed, we contrasted residual F_ST_ between antagonistic and non-antagonistic windows using Wilcoxon Rank-Sum tests.

### Linkage-based analyses of balancing selection

We examined the extent to which antagonistic haplotypes are selectively maintained by investigating whether antagonistic SNPs have unusually high linkage disequilibrium (LD) in the ZI population, the population that is most distant from LH_M_ and where levels of LD between antagonistic SNPs should be weakest in the absence of long-term balancing selection. Thus, for all SNPs situated within 1000bp of one another in ZI and which were also covered in LH_M_ (*i.e.*, SNPs which could be inferred to be either antagonistic or non-antagonistic), we calculated pairwise LD (r^2^) in PLINK. We then compared r^2^ values between pairs of antagonistic SNP and two control pairs: non-antagonistic pairs, and ‘mixed’ pairs (antagonistic/non-antagonistic). Comparing pairs of antagonistic SNPs to the mixed pairs allowed us to consider only SNPs located close to an antagonistic SNP, thus effectively controlling for possible non-random distributions of antagonistic pairs and non-antagonistic pairs with respect to genome-wide recombination rates.

To test for significant differences in LD between antagonistic pairs and the two control pairs, we modelled variation in r^2^ as a declining exponential function of chromosomal distance, and assessed differences in residual r^2^ (once distance was regressed out) using Wilcoxon Rank-Sum tests.

### Statistical software

All statistical analyses were carried out in RStudio (version 1.0.136^75^).

### Data availability

Phenotypic data will be deposited on Dryad prior to publication.

Population genomic data from LH_M_ is available at https://zenodo.org/record/159472. Population genomic data from the DGRP, ZI and SA is available at http://www.johnpool.net/genomes.html.

### Code availability

Analysis code is available on request.

## Acknowledgements

We are grateful to Aida Andrés for her suggestions on the analysis of balancing selection and to Laurent Keller, Andrew Pomiankowski and François Balloux for helpful comments on earlier versions of the manuscript. FR was funded by a London NERC DTP studentship (NERC grant NE/L002485/1), MSH was funded by a UCL IMPACT PhD studentship, EHM, TMP, IF and FCI by a European Research Council Grant (280632) and a Royal Society University Research Fellowship and MR and KF by a Natural Environment Research Council research grant (NE/G0189452/1).

## Author contributions

EHM, MR, KF, MSH and FR conceived the project; TMP, IF and EHM conducted phenotypic experiments; FR, MSH, MR and FCI analysed the data; FR, MR, MSH and KF wrote the manuscript.

